# A dynamic *WUSCHEL* / Layer 1 interplay directs shoot apical meristem formation during regeneration

**DOI:** 10.1101/2024.01.16.575947

**Authors:** Manoj Kumar, Dana Ayzenshtat, Hanita Zemach, Eduard Belausov, Leor Eshed Williams, Samuel Bocobza

## Abstract

*De novo* shoot apical meristem (SAM) organogenesis during regeneration in tissue culture has been investigated for several decades, but the precise mechanisms governing early-stage cell fate specification remain elusive. In contrast to SAM establishment during embryogenesis, *in vitro* SAM formation occurs without positional cues, and is characterized by spontaneous cellular patterning. Here, we have elucidated the initial stages of SAM organogenesis and the molecular mechanisms that orchestrate gene patterning to establish SAM homeostasis. We found that SAM organogenesis in tobacco calli initiates with protuberance formation followed by the formation of an intact L1 layer covering the nascent protuberance. Acquisition of L1 cellular identity is indispensable for de novo SAM formation, which also requires *WUSCHEL* (*WUS*) and the cellular capacity to direct anticlinal cell divisions. An intriguing finding is that *TONNEAU1* silencing prevents the exclusive occurrence of anticlinal divisions in the outermost layer of the protuberances and suppresses the acquisition of L1 cellular identity, ultimately impeding regeneration. This study exposes an intricate interplay between L1 and *WUS* expression and that any disruption in this interplay compromises shoot formation. It further provides a novel molecular framework for the characterization of *WUS*/L1 interplay-mediated shoot apical meristem formation during regeneration.

## Introduction

Plants exhibit a remarkable capacity to regenerate tissues, organs, or even an entire plant body. In nature, this process is used to recover from injuries or in asexual reproduction. For over 70 years, researchers have utilized this capacity in plant tissue culture systems to study cell differentiation and organ regeneration. Early studies demonstrated that organ formation in tissue culture systems relies on the interplay between two phytohormones, auxin and cytokinin (Skoog and Miller, 1957). The relative ratios of these hormones determine whether shoot, root, or callus formation is induced. Following studies explored the processes involved in shoot regeneration from callus in tobacco utilizing histological analyses revealed that the initial step is marked by the emergence of actively dividing cell regions in the section of the callus that faces the media (Thorpe and Murashige, 1968, 1970; Ross et al., 1973). The subsequent appearance of broad protuberances on the callus surface, composed of elongated parenchymatous cells, implies that these structures experience organogenesis to develop into meristemoids (E. Maeda and Thorpe, 1979). Studies on *Convolvulus arvensis* led to the conclusion that regeneration can be divided into three stages: competence, induction, and development (Christianson and Warnick, 1985). According to this model, competence refers to the acquisition of regenerative capacity, while induction denotes the initiation of regeneration, where the developmental fate of competent cells is determined. The third stage corresponds to organogenesis and to the development of the organ determined in the second stage. Despite a long history of research in this field, the precise sequential events and stages that govern the complex organogenesis of shoot meristem during regeneration remain unclear (Gary S. Hicks, 1994; Eshed Williams, 2021).

Shoot formation from the callus involves massive cellular reprogramming and *de novo* organization of adjacent cells to form a new shoot apical meristem (SAM). The *de novo* formation of a SAM entails a comparable level of patterning and cellular organization to that observed in SAM formation during embryogenesis (Mayer et al., 1998; Gordon et al., 2007). From an anatomical perspective, the SAM is composed of three tissue layers, namely L1, L2, and L3 (Satina et al., 1940), where cells within the L1 and L2 layers divide in an anticlinal manner and give rise to the epidermis and ground tissue, respectively. The cells in the L3 divide both anticlinally and periclinally and give rise to internal tissues. Furthermore, the SAM can be divided into three functional zones, namely the central, peripheral, and rib zones (Walles, 1991; Gifford and Corson, 1971). At the top of the SAM, the central zone contains stem cells that give rise to descendants, which are pushed to the peripheral or the rib zone. The WUSCHEL (WUS) transcription factor is required for stem cell specification and, thus, for the initiation and further maintenance of the SAM. The expression of the *WUS* gene within a specific group of cells situated beneath the stem cells delineates the organizing center within the central zone (Laux et al., 1996). The WUS proteins move from the organizing center to the meristem apex via the plasmodesmata (Yadav et al., 2011; Daum et al., 2014) and induce the expression of *CLAVATA3* (*CLV3*) by directly binding to its promoter (Perales et al., 2016). In turn, the CLV3 peptide represses *WUS* expression by interacting with the CLV receptors. This *WUS–CLV3* negative feedback circuit between the stem cells and the underlying organizing center plays a crucial role in maintaining SAM homeostasis (Schoof et al., 2000). Ectopic *WUS* expression is sufficient to induce somatic embryo formation or *de novo* SAM formation in Arabidopsis and many other species (Zuo et al., 2002; Gallois et al., 2004; Rashid et al., 2007; Negin et al., 2017). Consequently, multiple mechanisms and factors tightly control *WUS* expression within the meristem, while silencing it outside the meristem.

One question that has long captivated researchers is how gene patterning is maintained within the SAM despite cells being constantly displaced from their domains. During shoot regeneration from callus, an even more intriguing question arises: How is the necessary gene expression patterning established at the onset of SAM organogenesis in the absence of initial positional information? The advent of Next-generation sequencing technologies, innovations in imaging technologies, including live-imaging and computational modeling, and the development of numerous fluorescent reporters facilitated the understanding of the principles and mechanisms underlying patterning within the SAM and during shoot regeneration (Eshed Williams, 2021). One of the newly discovered mechanisms for patterning in the SAM is the formation of multiple morphogenes in the L1 layer of the meristem. Because L1 is the outermost layer and is constantly defining the boundaries of the organism, it functions as a reference tissue for patterning. In so doing, L1 acts as a source of signals such as microRNAs and active cytokinins that generate gradients toward the inner SAM layers thereby generating differential responses (Han et al., 2020; Chickarmane et al., 2012). For example, the *MIR394* displays L1-specific expression and confers stem cell competence by repressing the F box protein LEAF CURLING RESPONSIVENESS in the underlying cells to potentiate *AtWUS* (Knauer et al., 2013). Another example is the L1 layer-specific expression of *AtLOG4* (Chickarmane et al., 2012), which activates cytokinin nucleotides (Kuroha et al., 2009) while the *ARABIDOPSIS HISTIDINE KINASE (AHK)* cytokinin receptors are expressed in deeper layers (Gruel et al., 2016), which together suggests that L1 may generate a source of apically derived cytokinin in the SAM (Chickarmane et al., 2012) to promote *WUS* expression (Wang et al., 2017; Meng et al., 2017; Zhang et al., 2017; Zubo et al., 2017) and stabilize WUS proteins (Snipes et al., 2018).

In the SAM, the *ARABIDOPSIS THALIANA MERISTEM LAYER 1* (*ATML1*) transcription factor displays epidermis-specific expression and is commonly used as an L1 identity marker (Lu et al., 1996; Sessions et al., 1999) *ATML1* expression is initially detected during the two-apical-cell stage of embryo development, and by the 16-cell stage, it becomes restricted to the outermost epidermal cell layer (Iida et al., 2019). Being L1-specific, ATML1 patterns the concentration gradient of *HAIRY MERISTEM* (*HAM)* in the established SAMs via *MIR171* activation (Han et al., 2020).

In tissue culture systems, high levels of cytokinin can facilitate the induction of *WUS* expression and subsequently *de novo* SAM formation (Shin et al., 2020). In Arabidopsis, exogenous cytokinin promotes an auxin response through local auxin biosynthesis and polar auxin transport, resulting in the formation of an auxin ring within the callus where cytokinin biosynthesis is inhibited (Cheng et al., 2013). This precise distribution of cytokinin is part of a molecular mechanism that underlies auxin-cytokinin cross talk (Cheng et al., 2013) and is essential for *WUS* induction and further meristem formation (Gordon et al., 2009).

While the molecular mechanisms, which are involved in gene patterning and impart SAM homeostasis are well described, little is known about how patterning is established during *de novo* shoot meristem formation. In this study, we explore *de novo* shoot meristem formation. Using tobacco plants as a model, we identified the initial events that take place at the onset of SAM formation and found that the formation of an intact L1 layer is a key event toward SAM organogenesis. We also explored the complex interactions between *NtWUS* and *NtML1* expressions, which are critical components in initiating gene patterning in the newly formed meristem. This study offers a novel molecular framework that seeks to enhance our comprehension of WUS/L1- mediated SAM formation during regeneration.

## Results

### Histological and SEM Analyses of d*e novo* SAM formation in tobacco leaf explants

Tobacco leaf discs cultured on shoot-inducing media (SIM) develop callus bulges on the explant surroundings (i.e.at the wound sites) within a week. During the third week, shoots begin to emerge, and by the fourth week, leaf primordia become visible (**Fig. 1a**). To further characterize the processes of *de novo* SAM organogenesis, we conducted histological and scanning electron microscopy (SEM) analyses at three time points after SIM induction. On day 7, we detected foci of small cells with dense cytoplasm in proximity to the callus surface, as evidenced by the darker staining (**Fig. 1b**). On day 14, we noticed protuberances of about 100 μm protruding from the callus periphery, exhibiting an outermost cell layer formed by anticlinal cell divisions, typically found in the L1 layer of the SAM (Poethig, 1984). These protuberances appeared solely upon exposure to SIM (**Supplementary Fig. 1**) and might represent the initial stage of SAM organogenesis. From day 20, we observed fully developed meristems (**Fig. 1b**). Furthermore, SEM analysis of explants from day 14 revealed evident shoots with developed leaf primordia, which was confirmed by the presence of trichomes on their surface (**Supplementary Fig. 2a, 2b**).

**Fig. 1:**
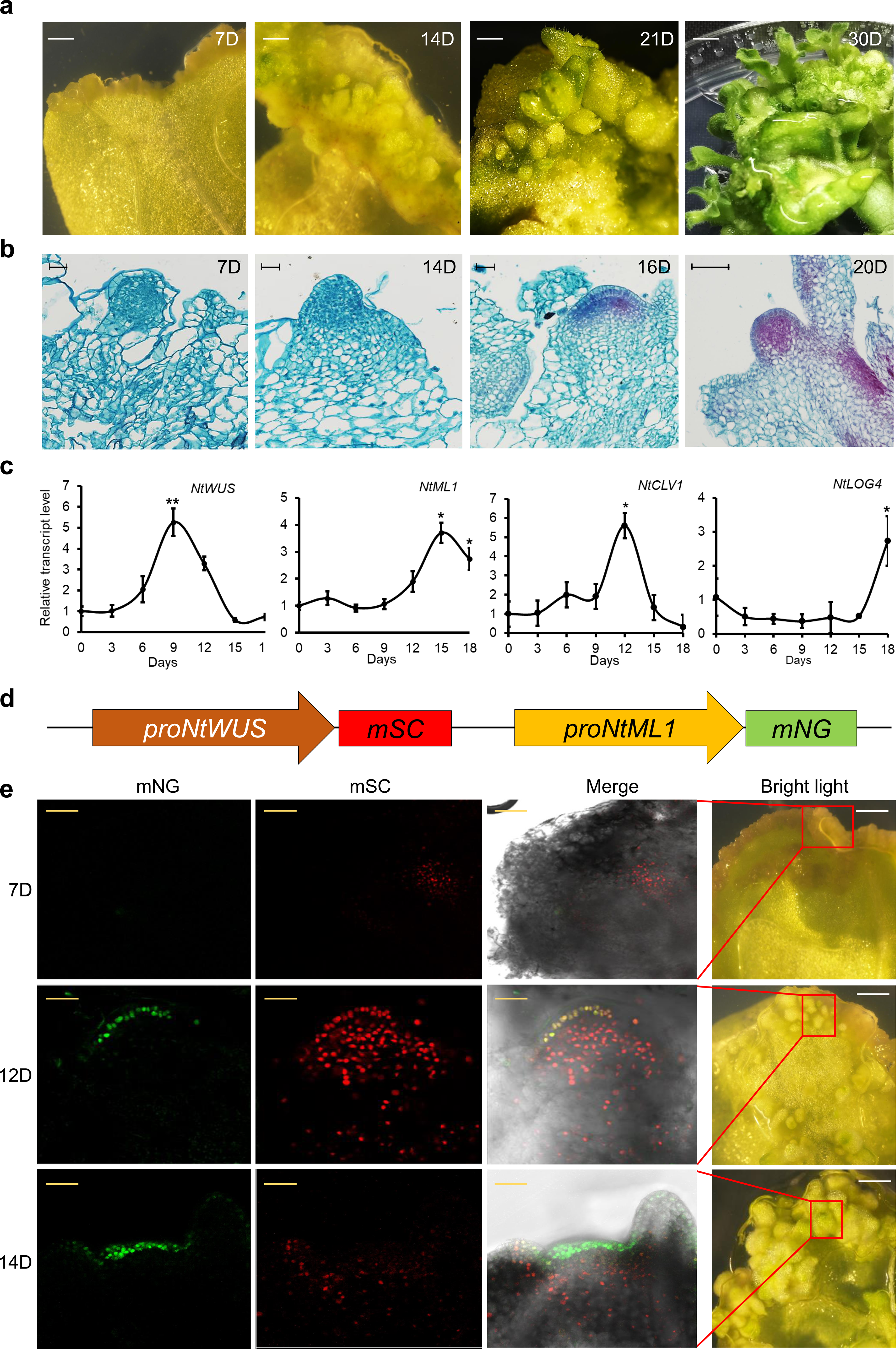
Induction of SAM formation and dynamics of *NtWUS* and *NtML1* expression patterns in tobacco leaf explants. **a.** Bright light images of tobacco leaf explants exposed to SIM for 7, 14, 21, and 30 days. Scale bar: 500 μm. **b.** Histological images of tobacco leaf explants exposed to SIM for 7, 14, 16, and 20 days. Scale bar: 50 μm. **c.** Relative transcript levels of *NtWUS, NtML1, NtCLV1*, and *NtLOG4* in wild-type tobacco explants exposed to SIM for 0, 3, 6, 9, 12, 15, and 18 days. The data represent the mean ± SD from three biological replicates (n=3) and were compared to t0 using the Student’s *t* test: \**P* < 0.05; \*\**P* < 0.01. **d.** Schematic representation of the T-DNAs built for this assay, consisting of the *mSC*, and *mNG* genes placed under the control of the *proNtWUS,* and pro*NtML1* respectively. **e.** Confocal images of transgenic tobacco leaf discs exposed to SIM for 7, 12, and 14 days. Scale bars: 20 μm (confocal), 1000 μm (bright light).

### Analysis of *NtWUS* and *NtML1* expression patterns during *de novo* SAM formation

To confirm that the protuberances observed at the early stages of shoot regeneration are newborn meristems and to characterize the identity of the cells in the epidermis-like layer of the protuberances, we investigated the expression of well-known meristematic genes in the developing protuberances. The *ATML1* gene, which encodes an epidermis-specific transcription factor, is involved in embryo formation and is commonly used as an L1 identity marker in the meristem (Lu et al., 1996). We identified the homologs of *WUS*, *ATML1*, *LOG4*, and *CLV1* from Arabidopsis in tobacco, which share 40%, 80%, 89%, and 84% amino acid sequence similarity, respectively (**Supplementary information**). Based on homology, we could not identify the homolog of *AtCLV3* in tobacco. Next, we exposed tobacco leaf disc explants to SIM and monitored the *NtWUS* (LOC107796712), *NtML1* (LOC107806765), *NtLOG4* (KU507075), and *NtCLV1* (LOC107817166) transcript levels in the periphery of the leaf discs at three-day intervals for 18 days using qPCR. *NtWUS* expression was initiated after day 3, peaked at day 9, followed by a gradual decrease until day 15, after which it remained low, while *NtCLV1* expression displayed a similar pattern but peaked at day 12, showing a 3 days delay. *NtML1* expression began after day 9 and gradually increased until day 15, while *NtLOG4* expression began after day 15 (**Fig. 1c**), suggesting L1 formation is concomitant to meristem establishment.

We next attempted to characterize *NtWUS* and *NtML1* expressions at the onset of SAM formation. For this, we employed a promoter-fluorescent marker approach to reveal the spatial pattern of expression of these genes. We generated transgenic plants harboring the two reporter genes *mScarlet-*I*-NLS* (*mSC*) and *mNeonGreen-NLS* (*mNG*) under the control of the *NtWUS* (*proNtWUS*) and *NtML1* (*proNtML1*) promoters, respectively (**Fig. 1d**). The L1/epidermis specificity of *proNtML1* was validated in a parallel assay using T1 plant shoots (**Supplementary Fig. 3**). Using qPCR, we analysed the relative transcript levels of *mSC* and *mNG* in these T1 transgenic plants at 0, 3, 6, 9, 12, 15, and 18 days after exposure to SIM media, and found they match the transcript levels of *NtWUS* and *NtML1*, respectively (**Supplementary Fig. 4**). Next, we exposed the leaf explants from selected T1 plants to SIM and followed the reporters’ fluorescences (**Fig. 1e**). On day 7, we observed the development of protuberances that displayed a weak *proNtWUS::mSC* signal, only in a small group of cells within inner layers, while no *proNtML1::mNG* signal was detected at this stage (**Fig. 1e and Supplementary Fig. 5**). On day 12, the mSC signal expanded throughout the protuberances and the mNG signal appeared only at their outer layer. At this stage, both markers showed signals in the outer layer. On days 14 and 21, the protuberances appeared to develop into meristem-like structures. The mSC signal became confined to the inner cell layers and was absent from the outer layer of the meristem, while an intense mNG signal remained on the outermost single-cell layer, suggesting that it might be the L1 layer of the newborn meristem (**Fig. 1e** and **Supplementary Fig. 5**). To confirm the specificity of the *NtWUS* promoter in this dynamic expression, we generated transgenic plants with a control construct containing the *proNtML1::mNG* and *proAtUBIQ::RFP*, which directs constitutive *RFP* expression (**Supplementary Fig. 6**). Conducting the same experiment, we observed overlapping signals of mNG and RFP in the outermost layer of the protuberances at all time points tested till day 21 (**Supplementary Fig. 6**).

In summary, these results suggest that the initial step of *de novo* SAM organogenesis *in vitro* is the formation of protuberances, in which *NtWUS* expression initiates from deeper cell layers and further expands across the developing protuberances. As the outermost cell layer is formed, coupled with the specific onset of *NtML1* expression at this layer, indicating an identity of epidermal cells, *NtWUS* expression becomes confined to the inner layers, suggesting the newly formed L1 may have a role in suppressing *NtWUS* expression in this layer.

### Alterations of L1 identity blocks early events of SAM formation

The L1 single-cell layer identity is dictated, at least partly, by the specific expression of the *ML1* gene (Sessions et al., 1999; Takada et al., 2013). To further evaluate the involvement of *NtML1* in *de novo* SAM formation, we used a *proNtML1*-directed RNAi approach to silence this gene and the *proSlUBIQ* to overexpress it ectopically, together with the *proNtML1::mNG* and the *proAtUBIQ::RFP* markers (**Fig. 2a**). We transformed tobacco leaf discs with these constructs and placed the discs on SIM. We validated the down- and up-regulation of *NtML1* expression in the RNAi and overexpression assays, respectively, by qPCR analysis on day 21 (**Fig. 2b**). Also, based on its homology (**Supplementary information**), we identified *NtCER5* (LOC107828914), a homolog of the direct target of *ATML1* in Arabidopsis (Takada et al., 2013), and found elevated *NtCER5* transcript levels in the *NtML1* overexpressing explants, thereby validating the functionality of *NtML1* overexpression (*NtML1*-OE). Remarkably, both RNAi and overexpression assays caused the formation of brownish ball-shaped callus aggregates from which merely a few shoots were formed, compared with the control assay, in which we observed the formation of green ball-shaped calli that further developed into numerous healthy shoots (**Fig. 2c,d; Supplementary Fig. 7** and **Supplementary Fig. 8**). SEM analyses of those leaf discs did not reveal SAM formation when *NtML1* was down-regulated or overexpressed (**Supplementary Fig. 7** and **Supplementary Fig. 8**), and histological analyses further revealed the absence of the L1-like layer in the protuberances upon *NtML1* down-regulation (**Fig. 2e**). Surprisingly, when we analysed *NtWUS* transcript levels by qPCR, we found that both *NtML1*-RNAi and *NtML1*-OE caused a significant ∼two-fold increase in *NtWUS* expression. Following the expression pattern of *NtML1* by confocal microscopy, we found that *NtML1* RNAi and overexpression resulted in the clear absence of *proNtML1::mNG* signal in the outermost cells of the protuberances, compared with the strong and specific signal that was observed in the control assay (**Fig. 2f, Supplementary Fig. 7** and **Supplementary Fig. 8**). The lack of *proNtML1::mNG* signal suggests that precise *NtML1* expression is required for the complete acquisition of L1 identity. These results suggest that the formation of new meristems requires the initial development of L1 in the outermost cell layer and the specific expression of *NtML1* within this layer.

**Fig. 2:**
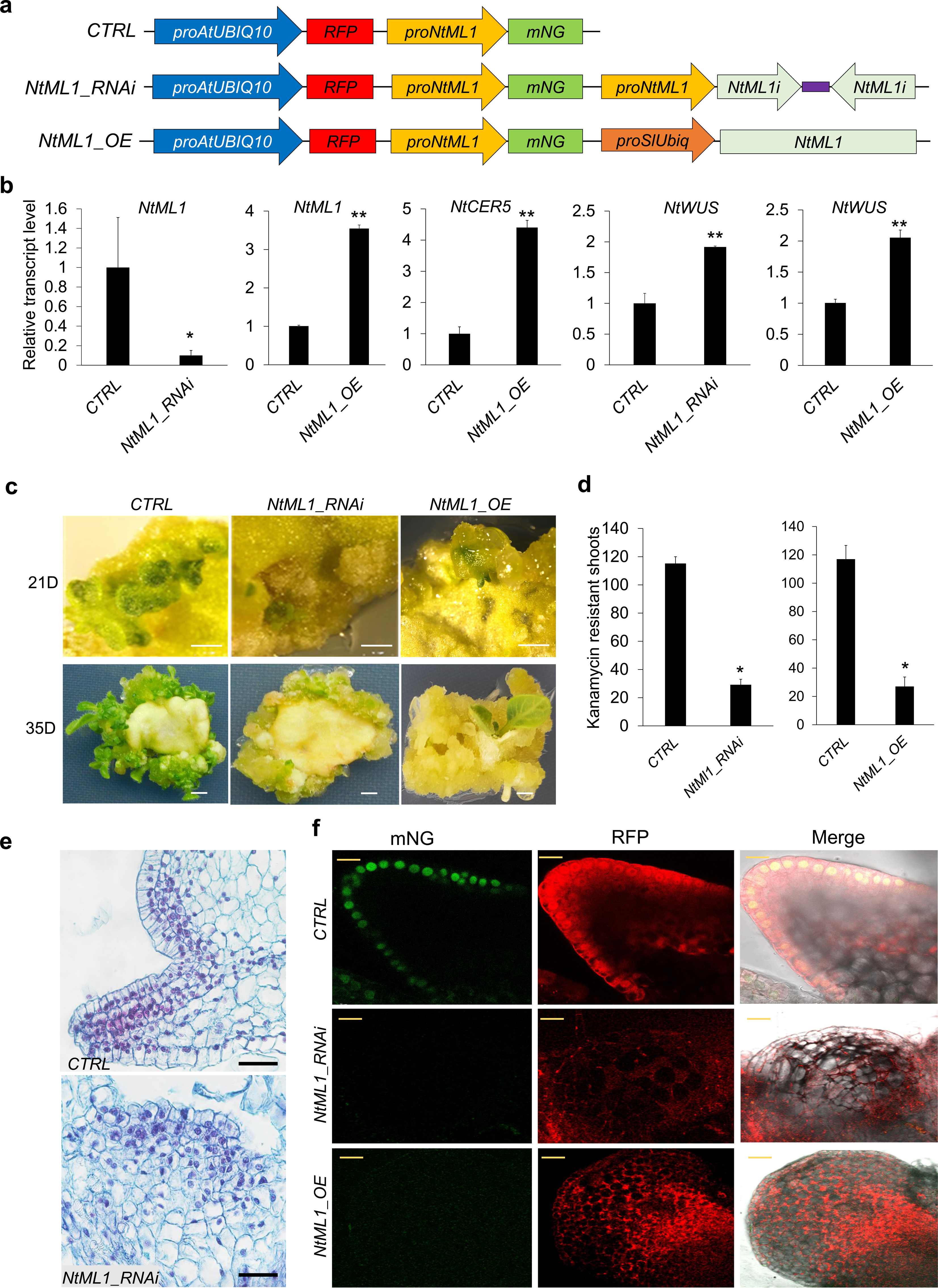
Alterations of L1 identity block SAM formation. **a.** Schematic representation of the T-DNAs built for this assay, consisting of *proNtML1*-directed RNAi designed to silence *NtML1* in L1 and *proSlUbiq*-directed *NtML1* overexpression cassettes, and the *mNG* and *RFP* marker genes placed under the control of the pro*NtML1* and *proAtUbiq*, respectively. **b.** Relative transcript levels of *NtML1*, *NtWUS*, and *NtCER5* in tobacco leaf discs transformed with the abovementioned T-DNAs and exposed to SIM for 21 days, n=3; Student’s *t* test: \**P* < 0.05; \*\**P* < 0.01. **c.** Bright light images of tobacco leaf discs transformed with the abovementioned T-DNAs and exposed to SIM for 21 and 35 days. Scale bars: 2000 μm (21D), 500 μm (35D). **d.** Bar graph depicting the number of developing shoots recorded 50 days after transformation (DAT) with the abovementioned T-DNAs, n=3; Student’s *t* test: \**P* < 0.05; \*\**P* < 0.01. **e.** Histological analysis of tobacco leaf discs transformed with the *NtML1*-RNAi and control constructs and pictured 21 DAT. Scale bar: 50 μm. **f.** Confocal imaging of tobacco explants transformed with the abovementioned T-DNAs and pictured 21 DAT. Scale bar: 20 μm.

### DTA-mediated L1 layer ablation blocks early events of SAM formation

To test the necessity of the putative epidermis layer of the protuberances for *de novo* SAM formation, we took the diphtheria toxin chain A (DTA) approach for genetic ablation (Czakó et al., 1992). This method enables the destruction of specific cells while maintaining the structural and functional integrity of the remaining tissue (Tsugeki and Fedoroff, 1999; Nilsson et al., 1998; Day et al., 1995). To specifically disrupt the L1 layer, we designed a construct that directs L1-specific *DTA* expression using *proNtML1*, while using a constitutive *DTA* expression construct as a control. Both constructs were fused to the *proNtML1::mNG* and the *proAtUBIQ::RFP* markers (**Fig. 3a**). Following Agrobacterium-mediated tobacco transformation, explants were cultured on SIM supplemented with kanamycin. Remarkably, both the *proNtML1*-derived and the ubiquitous expression of *DTA* drastically reduced the overall number of shoots obtained, whereas the control assay, which did not contain *DTA*, produced numerous viable shoots (**Fig. 3b** and **Supplementary Fig. 10**). Using SEM analysis, we did not observe meristem formation when DTA was used with either promoter (**Supplementary Fig. 9**). Next, we analysed these explants on day 21 after transformation using confocal microscopy and found that although the *proNtML1::DTA* produced protuberances, they were not covered by a single-cell layer of mNG-expressing cells. Instead, the *proNtML1::mNG* signal was disordered and appeared randomly in a few cells only, demonstrating that the L1 layer is altered (**Fig. 3c** and **Supplementary Fig. 9**). In explants transformed with *proSlUBIQ::DTA*, the *proNtML1::mNG* signal was blurred and not restricted to the nuclei, despite the use of a nuclear localization signal, suggesting *DTA* over-expression caused ectopic tissue degradation. In contrast, the outermost layer of the protuberances in the control assay (without *DTA*) exhibited a clear *proNtML1::mNG* signal (**Fig. 3c** and **Supplementary Fig. 9**).

**Fig. 3:**
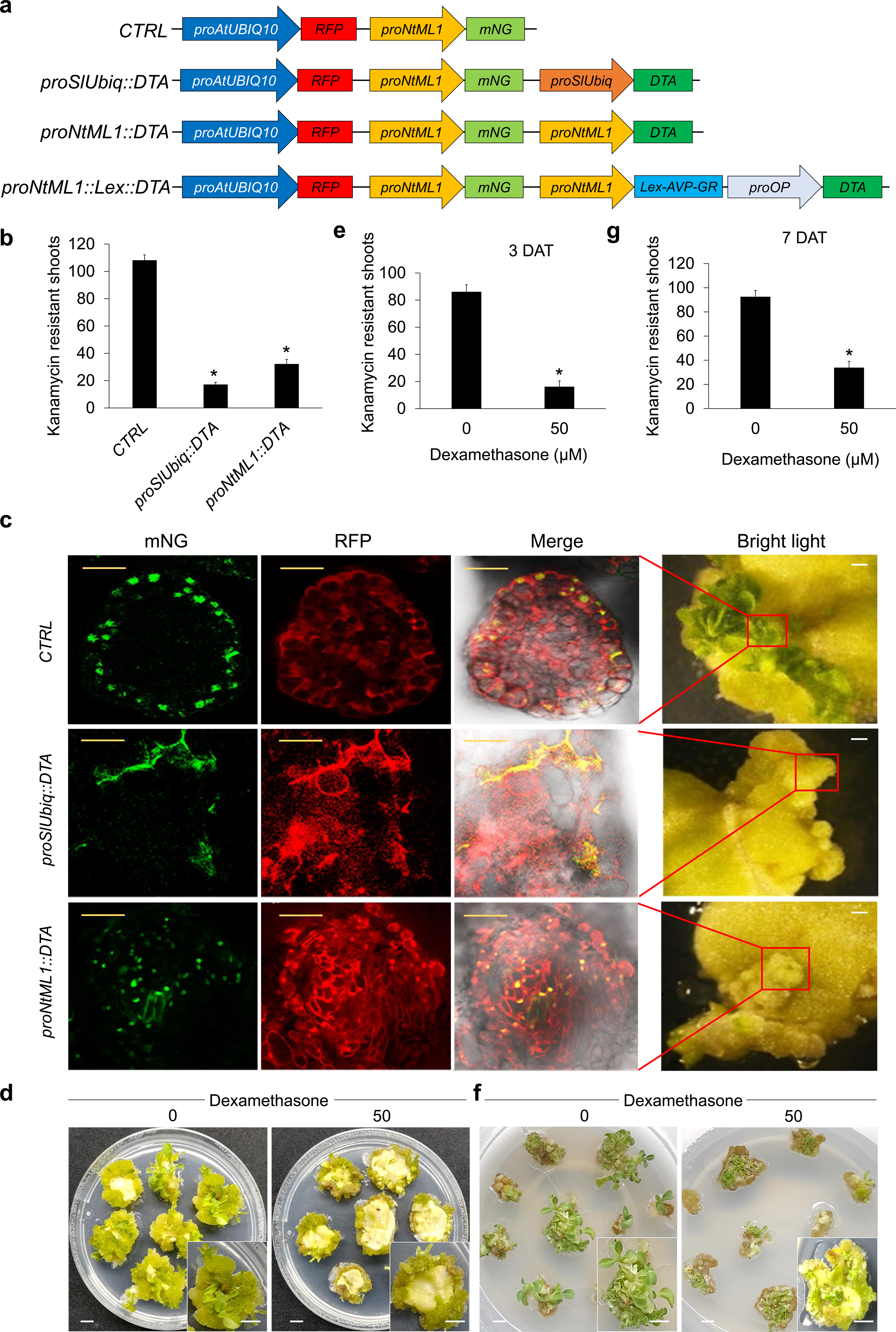
DTA-mediated L1 layer ablation blocks early events of SAM formation. **a.** Schematic representation of the T-DNAs built for this assay, consisting of *DTA* directed by the *proSlUbiq*, *proNtML1*, or under ML1-directed dexamethasone-induction, and of the *mNG*, and *RFP* genes placed under the control of the pro*NtML1* and *proAtUbiq*, respectively. **b.** Bar graph depicting the number of developing shoots recorded 50 DAT with the mentioned constructs, n=3; Student’s *t* test: \**P* < 0.05; \*\**P* < 0.01. **c.** Confocal imaging of tobacco explants transformed with the abovementioned T-DNAs and pictured 21 DAT. Scale bar: 20 μm. **d.** Bright light images of tobacco explants transformed with the inductive construct and exposed to Dex treatments (0 and 50 µM) 3 DAT and pictured 40 DAT. Scale bar: 500 μm. **e.** Bar graph depicting the number of developing shoots recorded 40 DAT while exposed to Dex treatments 3 DAT, n=3; Student’s *t* test: \**P* < 0.05; \*\**P* < 0.01. **f.** Bright light images of tobacco explants transformed with the inductive construct and exposed to Dex treatments (0 and 50 µM) 7 DAT and pictured 40 DAT. Scale bar: 500 μm. **g.** Bar graph depicting the number of developing shoots recorded 40 DAT while exposed to Dex treatments 7 DAT, n=3; Student’s *t* test: \**P* < 0.05; \*\**P* < 0.01.

Because L1-directed DTA is expected to disrupt the L1 layer, it is anticipated that *proNtML1::DTA* expression will cease in the absence of epidermal cell identity. Thus, we took advantage of the dexamethasone (Dex)-mediated gene expression system to direct the inducible *DTA* expression in the L1 layer. For this, we transformed tobacco leaf discs with a *proNtML1::LexAVPGR; pOP::DTA* construct (**Fig. 3a**) and cultured the tissue on SIM supplemented with kanamycin and Dex. To ensure that induced L1 destruction impairs early *de novo* SAM formation events, the transformed explants were exposed to Dex immediately (day 3) and seven days (day 7) after transformation. As expected, induced L1-directed *DTA* expression strongly restricted SAM formation, compared with the negative control in which Dex was not used (**Fig. 3d-g**).Together, these results supported the conclusion that eliminating *NtML1*-expressing cells suppresses the ability to establish a functional meristem.

### Disruption of cytokinin biosynthesis in L1 blocks SAM formation

The function of the L1 layer in the SAM includes the biosynthesis of active cytokinin, which is produced through L1-specific expression of *LOG4* and required to maintain gene patterning in Arabidopsis SAMs (Kuroha et al., 2009; Chickarmane et al., 2012). To evaluate the significance of this specific L1 function during *de novo* SAM formation, we took an RNAi approach to silence *NtLOG4* in an L1-specific manner, using *proNtML1*. This cassette was fused to the *proNtML1::mNG* and the *proAtUBIQ::RFP* markers (**Fig. 4a**) and used to transform tobacco leaf discs. The *NtLOG4* down-regulation was validated using qPCR (**Fig. 4b**). Similar to *NtML1* silencing, the *NtLOG4* down-regulation resulted in the formation of brownish, ball-shaped callus aggregates, from which only a few shoots formed, in comparison to the control assay (**Fig. 4c,d** and **Supplementary Fig. 10**). Using SEM microscopy, we could not detect SAM formation in *NtLOG4*- RNAi assays (**Supplementary Fig. 10**). In addition, *NtLOG4* down-regulation caused the disorganization of the cell layers in the protuberances, where the outer cells seemed fewer and larger cells, but still displayed *proNtML1::mNG* signals in their nucleus, implying that these cells differentiated (**Fig. 4e** and **Supplementary Fig. 10**). Using qPCR, we also observed a mild reduction in *NtML1* expression and an increase in *NtWUS* expression in the *NtLOG4* down-regulated explants (**Fig. 4f,g**). The reduction in *NtML1* expression could result from the fewer cells comprising the L1 layer. These results suggest that *NtLOG4* expression in the L1 layer is crucial for *de novo* meristem formation as this may be necessary to maintain a functional L1-cell layer. Additionally, it suggests that cytokinin biosynthesis is essential to preserve the integrity of the L1 layer.

**Fig. 4:**
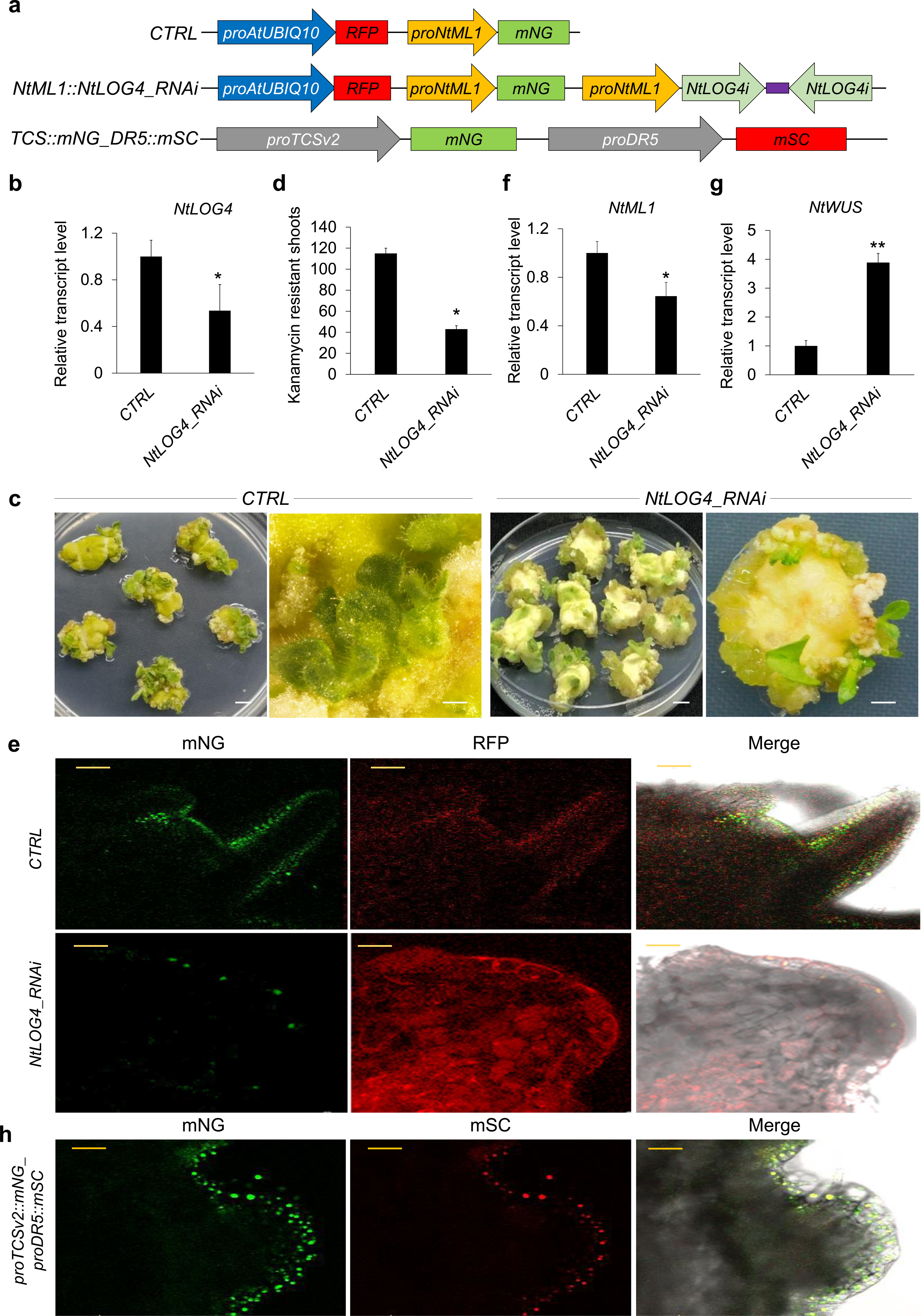
Alteration of cytokinin biosynthesis in L1 blocks SAM formation. **a.** Schematic representation of the T-DNAs built for this assay, consisting of a *proML1*-directed RNAi cassette designed to target *NtLOG4*, and the *mNG*, and *RFP* genes placed under the control of the pro*NtML1* and *proAtUbiq*, respectively, and of *mNG* and *mSC* directed by the *TCSv2* and *DR5* promoters, respectively. **b.** Relative transcript levels of *NtLOG4* in tobacco leaf discs transformed with the abovementioned T-DNAs and exposed to SIM for 21 days, n=3; Student’s *t* test: \**P* < 0.05; \*\**P* < 0.01. **c.** Bright light images of tobacco leaf discs transformed with the abovementioned T-DNAs and exposed to SIM for 21 days. **d.** Bar graph depicting the number of developing shoots recorded 50 DAT following transformation with the abovementioned T-DNAs, n=3; Student’s *t* test: \**P* < 0.05; \*\**P* < 0.01. **e.** Confocal imaging of tobacco explants transformed with the abovementioned T-DNAs and pictured 21 DAT. Scale bar: 20 μm. **f.** Relative transcript levels of *NtML1* in tobacco leaf discs transformed with the abovementioned T- DNAs and exposed to SIM for 21 days, n=3; Student’s *t* test: \**P* < 0.05; \*\**P* < 0.01. **g.** Relative transcript levels of *NtWUS* in tobacco leaf discs transformed with the abovementioned T-DNAs and exposed to SIM for 21 days, n=3; Student’s *t* test: \**P* < 0.05; \*\**P* < 0.01. **h.** Confocal imaging showing the mNG and mSC fluorescence directed by the *TCSv2* and *DR5* promoters, respectively, indicating cytokinin and auxin signaling at 7 DAT. Scale bar: 20 μm.

Furthermore, to determine the pattern of auxin and cytokinin signaling during early events of *de novo* SAM formation, we took advantage of the *DR5::mScarlet-I-NLS* and *TCSn::mNeonGreen-NLS* (Omary et al., 2022) (**Fig. 4h)**. In this assay, we cultured excised leaves from established T0 transgenic plants on SIM and followed the fluorescent signals at three time points. On day 5, small protuberances developed and displayed abundant mSC and NG fluorescent signals on their outer surface. On day 7, these signals were confined to the outer cell layer (**Fig. 4h**), and on day 14 we could observe clear fluorescence originating from the putative L1 layer that covered these protuberances (**Supplementary Fig. 11**). This result suggests auxin and cytokinin signalings are highly active in the outer cell layer of developing protuberances.

### *NtTON1* silencing restricts the anticlinal divisions in L1 and blocks SAM formation

Our results suggest that a functional L1 layer is crucial for shoot regeneration. Since the L1 layer is formed and characterized by its typical anticlinal plane divisions (Poethig, 1984; Kessler et al., 2006), we sought to disrupt the capacity of the L1-cells to direct anticlinal divisions and to investigate the impact of the disrupted L1 layer on shoot regeneration. To this end, we took advantage of the *TONNEAU1* (*TON1*) gene, which encodes a protein involved in the organization of cortical microtubules and in preprophase band (PPB) formation at the onset of mitosis to direct cell division plane positioning during cytokinesis (Spinner et al., 2010). The *ton1* mutants of Arabidopsis and *Physcomitrella patens* display abnormal patterns of cell division and position of division planes, and exhibit abnormalities in morphogenesis and cellular organization (Azimzadeh et al., 2008; Spinner et al., 2010). Since cells in the L1 layer require directed division planes, we tested the involvement of *NtTON1* in establishing a functional L1 layer and SAM. For this, we identified the *TON1* homolog of tobacco (LOC107818217), which shares 70.06% homology at the protein level with its Arabidopsis counterpart (**Supplementary information**). Next, we prepared RNAi constructs designed to target *NtTON1* in a ubiquitous and L1-specific manner using *proSlUbiq* and *proNtML1*, respectively. This cassette was fused to the *proNtML1::mNG* and the *proAtUBIQ::RFP* markers (**Fig. 5a**) and used to transform tobacco leaf discs. Silencing *NtTON1* with each construct resulted in the formation of protuberances that failed to form shoots (**Fig. 5b,c** and **Supplementary Fig. 12**). These protuberances exhibited clear *proAtUBIQ::RFP* signals, confirming successful plant transformation. However, no *proNtML1::mNG* signal was observed at 21 days post-transformation, indicating the absence of L1 cells (**Fig. 5d**). The *NtTON1*-RNAi lines showed aberrant division planes and alteration in cell shape, size, and number of nuclei within a cell, indicating aberrant cell divisions (**Fig. 5d**, and **Supplementary Fig. 12**). Remarkably, this result provides the first evidence of *NtTON1*’s involvement in cell division in tobacco, suggesting a functional similarity to the Arabidopsis *TON1*. These findings highlight the significance of anticlinal divisions in promoting L1 cellular identity and expose the involvement of *NtTON1* in directing cell plane orientation. Most importantly, these results indicate that proper L1 formation is a prerequisite for *de novo* SAM formation.

**Fig. 5:**
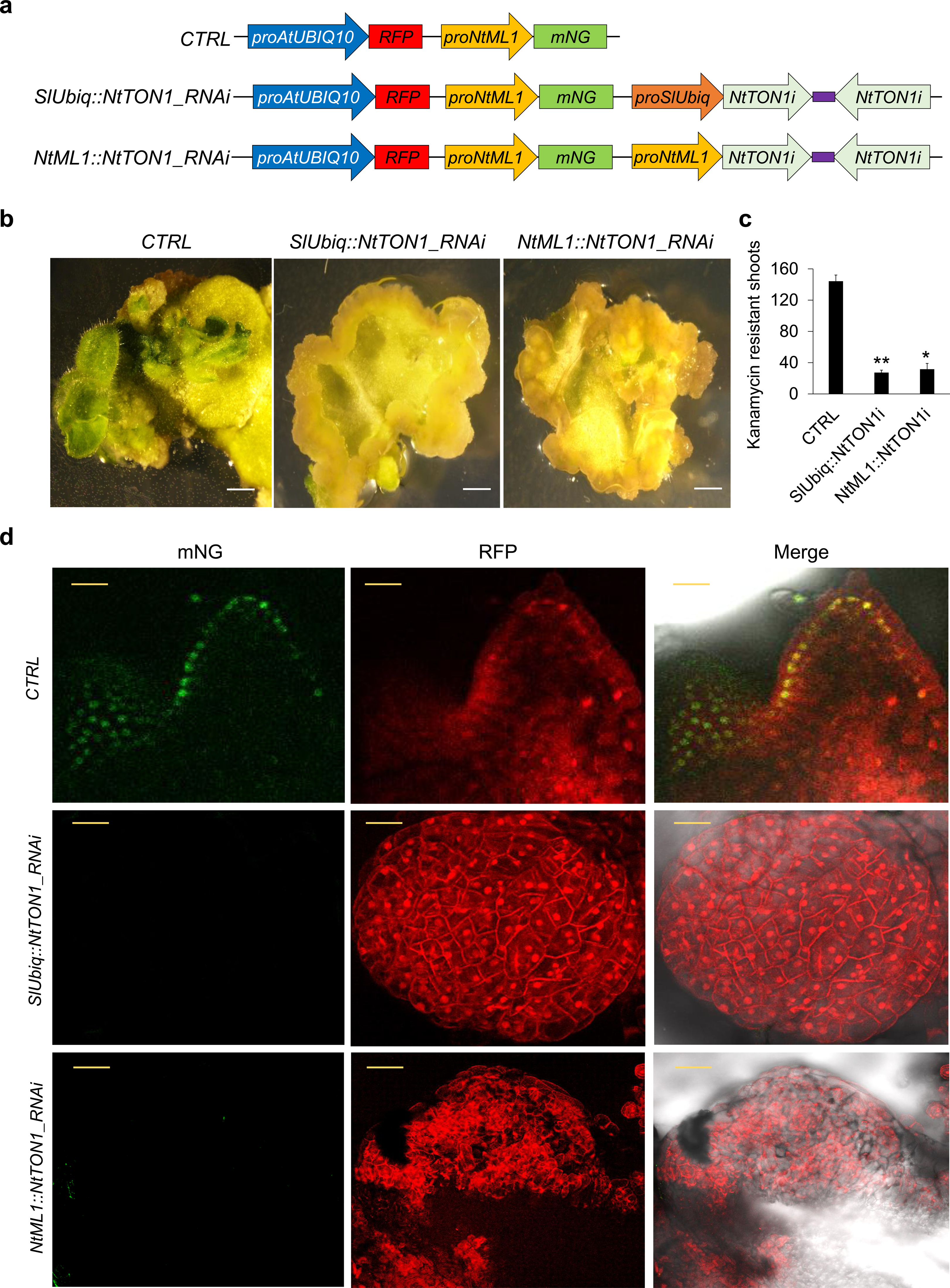
Silencing *NtTON1* blocks L1 and SAM formation. **a.** Schematic representation of the T-DNAs built for this assay, consisting of *proML1*- and *proSlUBIQ*-directed RNAi cassettes designed to target *NtTON1*, and the *mNG*, and *RFP* genes placed under the control of the pro*NtML1* and *proAtUbiq*, respectively. **b.** Bright light images of tobacco leaf discs transformed with the abovementioned T-DNAs and exposed to SIM for 21 days. Scale bar: 500 μm. **c.** Bar graph depicting the number of developing shoots recorded 50 DAT following transformation with the abovementioned T-DNAs, n=3; Student’s *t* test: \**P* < 0.05; \*\**P* < 0.01. **d.** Confocal imaging of tobacco explants transformed with the abovementioned T-DNAs and pictured 21 DAT. Scale bar: 20 μm.

### Alterations of genes involved in meristem maintenance block early events of SAM formation

Using DTA and RNAi approaches, we showed that the formation of a functional L1 layer is required for shoot regeneration. To further investigate *de novo* meristem organogenesis and the interdependency between meristem establishment and the L1 layer, we further silenced genes involved in meristem maintenance using RNAi. For this, we identified *NtWUS* (LOC107796712), *NtHAM* (LOC107824260), *NtPIN1* (LOC107830898), and *NtCLV1* (LOC107817166) (**Supplementary information**) and designed RNAi constructs to silence them. Each of these constructs was fused to the *proNtML1*::*mNG* and *AtUBIQ*::*RFP* fluorescent markers (**Fig. 6a**) and used to transform tobacco leaf discs. Down-regulation of *NtWUS*, *NtHAM*, *NtPIN1*, and *NtCLV1* led to a reduction in shoot formation and caused the formation of brownish ball-shaped callus aggregates that did not develop further, compared with the control assay (**Fig. 6b,c** and **Supplementary Fig. 13**). Silencing *NtHAM*, *NtPIN1*, and *NtCLV1* resulted in irregular fluorescent *proNtML1::mNG* signals, suggesting that proper SAM homeostasis is necessary for correct *NtML1* expression (**Supplementary Fig. 13**). However, the most intriguing phenotype was observed in the *NtWUS*-RNAi transformants, which displayed a complete absence of *proNtML1::mNG* fluorescence (**Fig. 6d**). Closer examination of the *NtWUS*-RNAi protuberances through histological analysis revealed their outer layer were disorganized and displayed both anticlinal and periclinal divisions (**Fig. 6g**), suggesting that *NtWUS* plays a crucial role in the L1 formation process. Quantification of *NtWUS* and *NtML1* transcript levels further supported this result by demonstrating a significant reduction in *NtWUS* and *NtML1* expression (**Fig. 6e,f**). This finding suggests that NtWUS is required to establish the cellular identity of the L1 layer, and may be necessary to initiate *NtML1* expression.

**Fig. 6:**
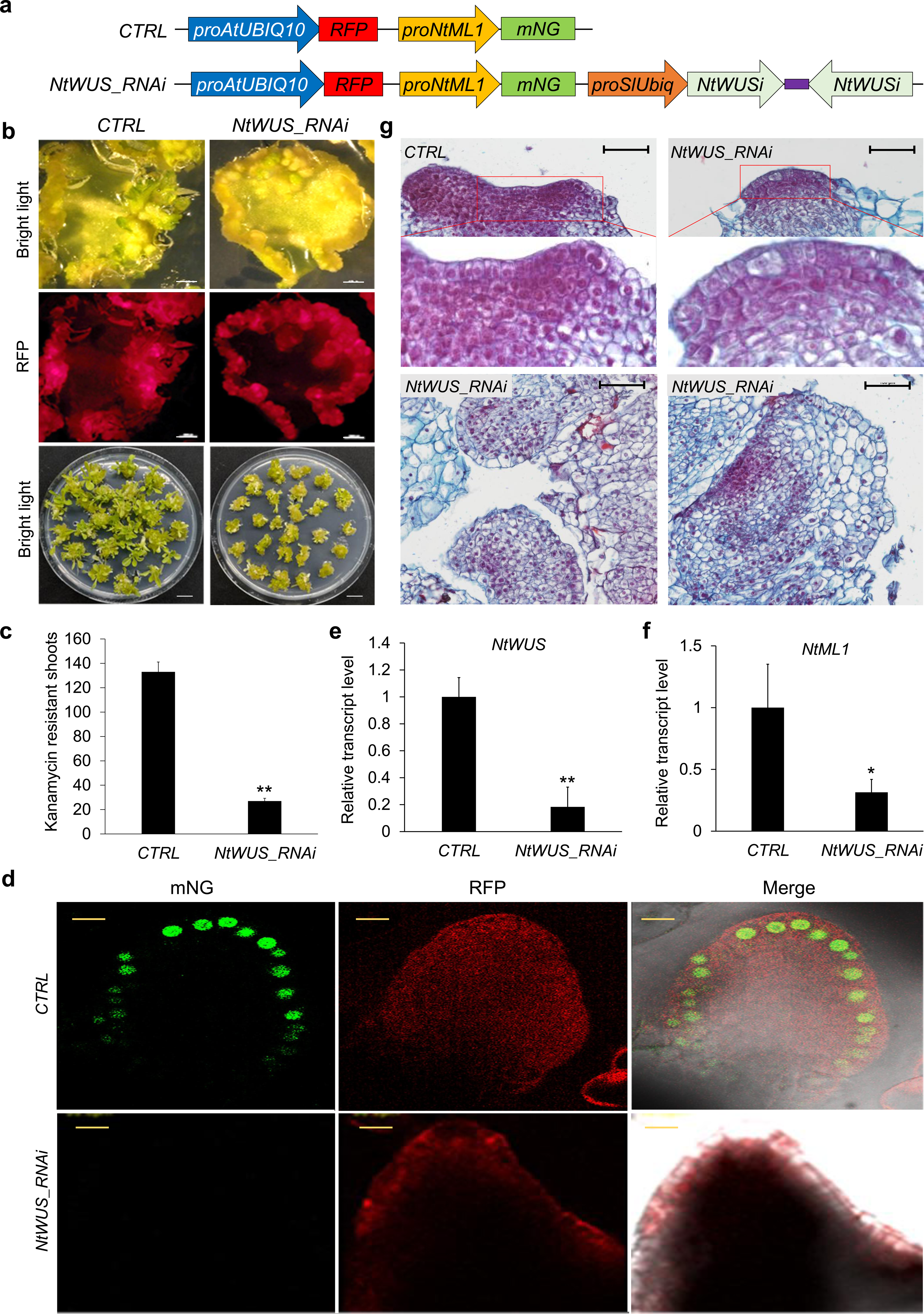
Alterations of meristem function block early events of SAM formation. **a.** Schematic representation of the T-DNAs built for this assay, consisting of *proSlUBIQ*-directed RNAi cassettes designed to target *NtWUS,* and of the *mNG* and *RFP* genes placed under the control of the pro*NtML1* and *proAtUbiq*, respectively. **b.** Bright light and fluorescence images of tobacco leaf discs transformed with the abovementioned T-DNAs and exposed to SIM for 21 days. Scale bar: 500 μm. Bright light images of explants on petri plates were taken 45 DAT. Scale bar: 1 cm. **c.** Bar graph depicting the number of developing shoots recorded 50 DAT following transformation with the abovementioned T-DNAs, n=3; Student’s *t* test: \**P* < 0.05; \*\**P* < 0.01. **d.** Confocal imaging of tobacco explants transformed with the abovementioned T-DNAs and pictured 21 DAT. Scale bar: 10 μm. **e.** Relative transcript levels of *NtWUS* in tobacco leaf discs transformed with the abovementioned T-DNAs and exposed to SIM for 21 days, n=3; Student’s *t* test: \**P* < 0.05; \*\**P* < 0.01. **f.** Relative transcript levels of *NtML1* in tobacco leaf discs transformed with the abovementioned T- DNAs and exposed to SIM for 21 days, n=3; Student’s *t* test: \**P* < 0.05; \*\**P* < 0.01. **g.** Histological images of tobacco explants transformed with the *NtWUS*-RNAi construct and pictured 20 DAT. Scale bar: 100 μm.

## Discussion

*De novo* organogenesis of shoot meristems from tobacco leaf discs begins with callus formation at the injury site and further develops without endogenous positional cues, except for the cells’ position relative to the media, and proximity to the original leaf tissue. This is in contrast to SAM establishment during embryogenesis, in which pre-patterning positional cues derived from the polar organization of the ovule play a crucial role in instructing the initial cellular patterning of the embryo (Capron et al., 2009). Considering that the protoderm is the first tissue formed during embryogenesis, its formation was suggested to be a prerequisite for the correct progression of embryogenesis (Ueda and Kurihara, 2021). In this study, we show that SAM organogenesis from leaf discs involves two key aspects: shaping the distinctive dome-like structure of the SAM, and initiating the patterning of gene expression required for meristem formation and maintenance. We also identified the establishment of an intact L1 layer as a key event toward SAM formation. We further provided evidence for the importance of L1 layer formation and the correct *NtML1* expression for shoot regeneration through DTA-mediated L1 ablation, *NtML1* silencing and overexpression, and *NtTON1* silencing. Most intriguing, we found that reducing *NtTON1* expression disrupts the ability to properly direct cell division planes, which impairs the exclusive occurrence of anticlinal divisions in the outermost layer of the protrusions. Consequently, this hinders the formation of the L1 layer and the acquisition of L1 cellular identity, resulting in impaired regeneration. We also analysed the dynamics between *NtWUS* and *NtML1* and provided evidence that any disruption in this interplay compromises the capacity to regenerate shoots.

### Dynamics of *NtWUS* and *NtML1* interaction at the onset of SAM organogenesis

Studies on shoot regeneration from Arabidopsis roots suggest that several rounds of cell divisions in the presence of cytokinin are necessary to acquire the competence to express *WUS*, which is required to initiate SAM formation (Shemer et al., 2015; Che et al., 2007). Similarly, when we expose tobacco leaf discs to cytokinin, *NtWUS* expression initiates at an early stage of cell proliferation (day 7), as evidenced by both qPCR and fluorescent markers. Next, *NtWUS* expands throughout the protuberances (day 12), and upon the onset of *NtML1* expression, specifically in the outermost layer, *NtWUS* becomes excluded from the top layers of the developing meristem (day 14). The presence of an ML1 *cis* binding site (L1 box (Takada and Jürgens, 2007)) in the *NtWUS* promoter (**Supplementary information**) and the significant up-regulation of *NtWUS* expression in *NtML1-*RNAi protuberances suggest that NtML1 may directly restrict *NtWUS* expression in the L1 layer. In Arabidopsis, the *PROTODERMAL FACTOR2* (*PDF2*) is an *ATML1* paralog and these genes function redundantly in the epidermis (Ogawa et al., 2015). Our result is consistent with the ectopic formation of adventitious shoots on the *atml1-3/+ pdf2-4* Arabidopsis mutant cotyledons (Ogawa et al., 2015), suggesting ectopic *WUS* expression. In light of our results, the observation of Ogawa et al suggests that in the absence of functional ATML1 and PDF2, the repression on *WUS* is released. However, assuming that NtML1 directly represses *NtWUS* in L1, we would expect to observe *NtWUS* down-regulation when we overexpress *NtML1*, but the opposite was obtained, suggesting an indirect regulation. Thus, we suggest that the lack of L1 in *NtML1*-OE protuberances indirectly causes *NtWUS* up-regulation. Therefore, L1 activity, rather than *ML1* expression, may be the factor that restricts *WUS* expression. The absence of L1 in *NtML1*-OE protuberances can be explained by the disruption of position-dependent cell differentiation mechanisms, which typically enable neighboring cells to adopt different fates (Hung et al., 1998). In *NtML1*-OE protuberances, the presence of *NtML1* in all cells impedes the acquisition of L1 cellular identity. Since L1 is a source of signals that limit *NtWUS* expression and is absent in *NtML1*-OE, *NtWUS* is up-regulated in such protuberances. This is also consistent with the up-regulation of *WUS* obtained in the *NtLOG4*-RNAi protuberance, where *NtML1* expression was down-regulated together with *NtWUS* up-regulation. Taken together, these results suggest that the overexpression of *NtML1* impedes the acquisition of L1 identity.

Conversely, the sequential pattern of gene expression, where *NtWUS* is expressed first (on day 7) followed by *NtML1* (on day 12), alongside the decrease in *NtML1* expression in the *NtWUS*- RNAi protuberances, suggests that WUS plays a role in promoting *ML1* expression. This could occur through direct regulation by means of binding the *NtML1* promoter and activating its expression, or indirectly by, for example, promoting the development of L1, which in turn may promote *NtML1* expression. Previous studies support both possibilities. For the first suggestion, analysis of the *ATML1* promoter has revealed that a 101-bp fragment upstream to the transcription start site, which encompasses both an L1 box and a WUS-binding site, is sufficient for the specific expression of *ATML1* in the outermost cell layer (Takada and Jürgens, 2007). Two WUS-binding sites (Takada and Jürgens, 2007) and one L1 box (Han et al., 2020) were found in the *NtML1* promoter too (**Supplementary information**). For the second possibility, surgical removal of the epidermal cells to expose the mesophyll cells, which then adopt the outermost position, led to the activation of *ATML1* expression in these cells (Iida et al., 2023). This implies that surface-positional cues play a role in promoting *ATML1* expression in the outermost cells. The reduced accumulation of *ATML1* only in the inner daughter cells of the epidermis after sporadic periclinal division in the L1 layer (Iida et al., 2019), further supports this notion. Our analysis of protuberances that were formed in *NtWUS*-RNAi revealed a disrupted outermost cell layer exhibiting both anticlinal and periclinal cell divisions and showing no *proNtML1::mNG* signal, suggesting *NtWUS* is required to promote anticlinal divisions, acquisition of L1 cellular identity, and *NtML1* expression.

Nevertheless, our analysis cannot establish a cause-and-effect relationship between these two observations: Does a reduction in *NtWUS* lead to a direct decrease in *NtML1* expression, or alternatively, does the reduction in *NtWUS* expression compromise other processes such as anticlinal divisions, leading in turn to an impaired L1 layer formation and to *NtML1* misexpression? The fascinating finding that silencing *NtTON1* alters the exclusive occurrence of anticlinal divisions on the surface of the protuberances, thereby disturbing the formation of the L1 layer and leading to the loss of *proNtML1::mNG* signal, provides support for the idea that anticlinal divisions are essential for the acquisition of L1 cellular identity and for *NtML1* expression.

### L1 formation is a key event toward SAM organogenesis

In a functional SAM, the L1 layer provides major positional cues to pattern gene expression domains that are essential for SAM maintenance (Chickarmane et al., 2012; Eshed Williams, 2021). Our analyses suggest that the development of an outermost single cell layer, which eventually develops into the L1 layer, is a crucial event in initiating SAM *de novo* organogenesis, and that this process occurs before gene patterning takes place in the SAM. The L1 layer plays a substantial role in maintaining gene patterning in the active SAM. It is involved in regulating *WUS* expression through the production of CLV3 (Clark et al., 1995), setting the *HAM* concentration gradient via *miR171* activity (Han et al., 2020; Zhou et al., 2018; Hendelman et al., 2016), and exclusively expressing the *LOG4* gene (Chickarmane et al., 2012). Being L1-associated, these signals may only be functional after L1 forms and becomes active. Nevertheless, the precise trigger for L1 formation remains unknown. The formation of the outermost cell layer occurs through a sequence of anticlinal cell divisions, which requires directed cell divisions. It is widely accepted that division orientation follows the rule of setting the division plane at the shortest possible path. However, in many instances, mechanical stress and auxin signaling instruct the division plane (Serra and Robinson, 2020; Yeoman and Brown, 1971). One of the mechanisms that restrict ML1 activity to L1 was shown to involve mechanical signaling (Iida et al., 2023), but the process in which mechano-transduction promotes L1 differentiation remains to be explored. Based on our histology analysis, before the outermost cell layer can be detected, intense cell divisions occur on the callus edge interior to the surface, generating protuberances (observed as many small cells with dense cytoplasm). This suggests that collective turgor pressure from highly proliferating cells may generate an ‘inner pressure,’ which in turn may direct the tensile stress in the cell wall of the outermost cells (**Fig. 7**). A recent study on the meristem boundary revealed that divisions align with the direction of maximal tensile stress, thereby demonstrating the role of mechanical forces as instructive signals in determining the cell plane of division (Louveaux et al., 2016; Robinson, 2021), and suggesting that cell wall tension is the most fundamental determinant of the cell division plane orientation (Louveaux et al., 2016; Guérin et al., 2016). Since cells are glued together, strong tensional stresses exist on the outer layer of a crease (Hamant et al., 2008). This tensile force may also act on the outer surface of the dome-shaped protuberances observed here during *de novo* SAM organogenesis (**Fig. 7**). In addition, auxin plays an important role in increasing local cell wall loosening (Trinh et al., 2021), consequently affecting the mechanical forces exerted on it (Majda and Robert, 2018). Therefore, the accumulation of auxin observed at the surface of the protuberance, as indicated by *DR5::mSC*, may also play a role in altering the mechanical forces acting on the outermost cells, thus contributing to the directed cell divisions at the protuberance surface.

**Fig. 7:**
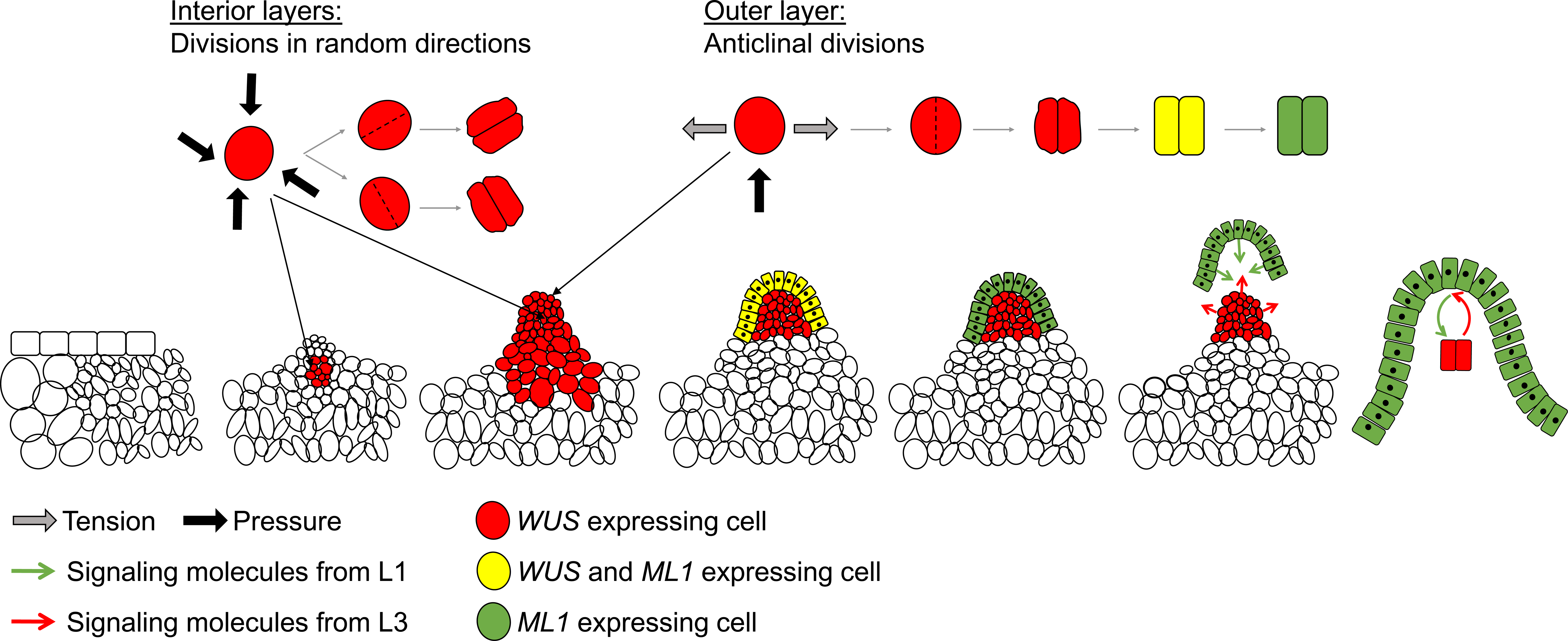
Schematic model depicting SAM development in tissue culture. The excised explant is depicted on the left. Callus cells first proliferate and further form a protuberance with restricted *WUS* expression at the center. Inside the protuberance, pressure is exerted on the cells and causes periclinal divisions, while tension applies to the outer cells and causes anticlinal divisions (Hamant et al., 2008). *WUS* expression further expands throughout the protuberance and *ML1* expression begins on its outer layer. Next, *WUS* expression becomes restricted to the center of the protuberance while *ML1* expression remains in L1. At last, L1 and L3 are established an secrete characteristic genetic information, yielding a functional SAM.

The significance of directed cell division to the formation of the L1 layer, and thus for *de novo* SAM organogenesis, becomes evident through the reduction of *NtTON1* expression by RNAi. In Arabidopsis, TON1 binds the preprophase band (PPB) during mitosis to indicate the future position of the cell plate, which defines the division plane (Drevensek et al., 2012; Van Damme et al., 2007). Accordingly, the Arabidopsis *ton1* mutant displays mis-orientated cell division (Azimzadeh et al., 2008). We found that reduced *NtTON1* expression in L1 allowed protuberance formation but did not allow the occurrence of anticlinal divisions and the *proNtML1::mNG* signal. Since the L1 cellular identity is dictated, at least partly, by the specific expression of the *ML1* gene in this single-cell layer (Sessions et al., 1999; Takada et al., 2013), this suggests that altered anticlinal divisions in the outer cell layer limit the complete acquisition of L1 identity. To our knowledge, this is the first evidence of the involvement of TON1 in directing the anticlinal division planes required for L1 formation and thus for *de novo* SAM formation.

Maintaining the L1 identity is crucial for further gene patterning in the newly formed meristem. In the SAM, the cytokinin promotes cell division through the regulation of D-type cyclin (Riou-Khamlichi et al., 1999). Here, we observed that reducing *NtLOG4* expression levels by RNAi resulted in the formation of larger cells in the L1 layer. It might be that in *NtLOG4*-RNAi assays, the outermost cells differentiate without active cytokinin to drive mitotic divisions, supported by the presence of trichomes on L1 (**Supplementary Fig. 10**). Given that the mechanism of gene patterning through gradients depends on morphogen mobility (Eshed Williams, 2021), it’s plausible that an increase in cell size impedes the capacity to establish a proper gradient, thereby interfering with establishing patterning during *de novo* organogenesis. The significant up-regulation of *WUS* in *NtLOG4*-RNAi assays may also be the result of improper gradients alongside the interference with the multiple spatial feedback loops between cytokinin and WUS (Gordon et al., 2009; To et al., 2004; Zhao et al., 2010).

## Conclusion

In this study, we intended to elucidate the intricate molecular mechanisms that govern SAM formation during shoot regeneration. We find SAM formation begins with the emergence of a distinctive dome-shaped protuberance, which subsequently becomes covered by an L1 cell layer. Due to its surface proximity, the outermost layer of the developing meristem may serve as a reference layer to produce positional cues for the precise patterning of gene expression domains. We examined the significance of this outermost cell layer in shoot regeneration at multiple levels: interfering with the cell layer formation (*NtTON1* down-regulation), destroying the L1 cells (using DTA), interfering with the acquisition of L1 cellular identity (altering *NtML1* expression), and deterring one of its functions (*NtLOG4* down-regulation). In each case, we observed remarkable alterations in both L1 formation and *ML1* expression, subsequently suppressing SAM formation. This compelling evidence underscores the indispensable role of the cellular identity and functions of the outermost layer in the *de novo* shoot formation process. Notably, while hormones and microRNAs have long been recognized as contributors to this process, the precise nature of the signals emanating from the L1 layer remains to be fully characterized. Most intriguing, we expose the critical role of *WUS* expression in orchestrating anticlinal mitotic divisions, which in turn are vital for L1 formation and *ML1* expression. On the other hand, the L1 formation also restricts *WUS* expression. Thus, the expression of *WUS* and *ML1* seems interconnected and any disruption in this interplay compromises the capacity to form shoots. Collectively, our findings establish the pivotal importance of establishing an active L1 layer in *de novo* SAM formation and maintenance. In summary, this study presents a novel and comprehensive molecular framework that sheds light on the interplay between *WUS* and L1 in SAM organogenesis during regeneration (**Fig 7**).

## Methods

### Cloning methods

All plasmids used in this study were assembled using the GoldenBraid cloning system (Sarrion-Perdigones et al., 2013). The *Neomycin Phosphotransferase-II* (*NPT-II*) gene was used as a selection marker for kanamycin resistance in all our constructs. The promoters such as *NtML1, NtWUS, SlUbiq, AtUbiq,* and *ZmUbiq* used here were cloned into entry vectors (pUPD; GoldenBraid) and further used to generate expression cassettes that were combined to produce multipartite assemblies, as described in the GoldenBraid recommendations. A 2041 bps region including *NtML1* promoter sequence and its 5’ untranslated region (UTR), and a 1738 bps region including *NtWUS* promoter sequence and its 5’ UTR were amplified by polymerase chain reaction (PCR) using primers sequences given in the **Supplementary Table 2**. These PCR products were cloned into an intermediate vector pUPD2 and sequencing was carried out to confirm the presence of the desired sequences. Furthermore, the *NeonGreen-NLS* (*mNG*) and the *mScarlet-I-NLS* (mSC) genes were fused to the *proNtML1* and to the *proNtWUS*, respectively, to generate reporter expression cassettes.

To construct the plasmids designed for RNAi, sense and antisense PCR products corresponding to the targeted genes including *NtCLV1*, *NtWUS, NtPIN1, NtHAM1, NtLOG4,* and *NtML1* were generated with Bsa1 restriction sites that left compatible sticky ends on both extremities (**Supplementary Table 2**), and were ligated with an intron fragment and either *proNtML1* or *proSlUbiq* into the alpha vector as described previously (Kumar et al., 2022). The resulting RNAi cassettes were then fused with *proNtML1::NeonGreen-NLS* and *proAtUbiq::RFP* cassettes.

### *Agrobacterium* transformation and preparation

The binary vectors generated in the present study were transformed to *Agrobacterium tumefaciens* strain EHA105 (Hood et al., 1993) using the freeze-thaw method (Höfgen and Willmitzer, 1988). Transformants were grown and selected on Luria broth (LB) media containing 10 mg/l rifampicin and 50 mg/l kanamycin (for p3alpha1 and p3alpha2 constructs) or 250 mg/l spectinomycin (for p3omega1 and p3omega2 constructs) at 28 °C until colonies appeared. Single colonies were picked and grown in liquid LB media at 28 °C and 200 rpm. The resulting suspension cultures were washed once in the infiltration medium consisting of 50 mM MES, 20 mM trisodium orthophosphate, 5 mg/l D-glucose, and 200 μM acetosyringone, and further resuspended in this media, which was adjusted to a final OD600 of 0.4.

Seeds of tobacco (*Nicotiana tabaccum*) plants were surface sterilized using chlorine gas sterilization procedure (Lindsey et al., 2017). For this, 150 ml of 6% sodium hypochlorite and 7 ml of 37% HCl were taken in a beaker and kept with the microfuge tube containing the tobacco seeds inside the dessicator in the chemical hood for 2 hours. Seeds were germinated and further grown on the germination medium. Plants’ nodes were separated and subcultured every 4 weeks. For morphogenic assays, leaf discs (1 cm) were excised from these plants’ leaves and were placed on the shoot-inducing medium (SIM), and the root-inducing medium (RIM; **Supplementary Table 1**). For stable transformations, twenty leaf discs were excised from 4-week-old plants grown *in vitro*.

The discs were incubated for 10 min with *Agrobacterium* containing the relevant plasmids. Then, the discs were dried on sterile filter papers and placed onto SIM, which consisted of MS (Murashige and Skoog, 1962) medium including vitamins supplemented with 3% sucrose, 3 mg/l 6- Benzylaminopurine (BA), 0.5 mg/l 1-Naphthalene acetic acid, and 8 g/l plant agar, pH = 5.8, for 48 h in the dark at 24 ± 2 °C. The explants were then transferred to selection media, which consisted of SIM, supplemented with 200 mg/l kanamycin and 250 mg/l timentin. The discs were transferred to new media at regular intervals of 10–12 days. After 5 weeks, the green elongated shoots were transferred into boxes containing growth media, which consisted of MS basal salt with Gamborg B5 vitamins, 3% sucrose, 8 g/l plant agar, pH = 5.8, 200 mg/l kanamycin, and 250 mg/l timentin to induce rooting. Green and healthy-looking shoots that had developed roots were transferred to pots filled with soil and further grown in our growth room. All the reagents and chemicals used for plant tissue culture purposes were purchased from Duchefa Biochemie, Netherlands.

### Histology

Leaf discs were placed on SIM for 0, 7, 14, and 20 days, and were collected and stored in FAA solution, which consisted of 10% formaldehyde, 5% glacial acetic acid, 52% absolute ethanol (v/v). Histochemical analyses were then performed as described previously (Singh et al., 2019). Briefly, samples were first dehydrated using ethanol dilution series, then embedded in paraffin wax before sectioning. Sections (12 μm) were prepared by using a microtome (Leica RM2245, Leica Biosystems, Nussloch, Germany), and were deparaffinized using a histoclear solution, before rehydration with ethanol dilution series. These processed sections were used for histochemical staining using 1 % (w/v) safranin and 0.2 % (w/v) fast green staining to observe the vascular system and lignin deposition. Microscopic examinations were performed using a light microscope (ECLIPSE Ni-E, Nikon) and images were taken with a Nikon DS-Fi1 digital camera.

### Scanning Electron Microscopy

Leaf discs were placed on SIM (**Supplementary Table 1**) and were collected at indicated days. Samples were fixated using the methanol-ethanol fixation method (Talbot and White, 2013). Briefly, samples were soaked in methanol for 10 min, then soaked in absolute ethanol for 30 minutes, further dehydrated overnight in absolute ethanol, and then stored at 4°C until used. Critical point dry (CPD) was conducted following the manufacturer’s recommendations (K850, QUORUM). Finally, tissue was mounted on an SEM stub, coated with palladium metal, and observed using a versatile benchtop JEOL (JCM-6000) scanning electron microscope.

### Measurement of RFP fluorescence in tobacco leaves

In the stable transformation assays, RFP fluorescence was observed in the transformed calli using fluorescent microscopy 21 days after transformation (DAT). This was performed using a fluorescent binocular (SMZ25 Stereomicroscope, Nikon instruments), in which settings were adjusted to 1 × magnification, 100 ms exposure, and 2.8 gain.

### Confocal Laser Scanning Microscopy

To detect the fluorescences of RFP, mNeonGreen (mNG), and mScarlet-I (mSC) proteins, manually dissected calli/apices were collected from *in vitro* transformed tobacco transformants. For confocal laser scanning an Olympus IX81/FV500 (Japan) laser-scanning microscope was used to observe fluorescently labeled cells with the following filter sets: for eGFP and mNG, 488 nm excitation, and BA505–525 were used; for RFP and mSC, 561 nm excitation and BA570-610 were used. The objective used was UPlanApo 20x/0.7 or PlanApo 40x/0.9 water (WLSM). When mNG and RFP or mSC were detected in the same sample, dichroic mirror 488/561 was used. We also performed sequential fluorescence acquisition when more than one fluorescent marker was monitored.

### Dexamethasone treatment and morphological observations

Transformed leaf disc explants of tobacco were co-cultivated with Agrobacterium on SIM for 3 days. They were then transferred to the selection medium supplemented with 200 mg/l kanamycin. The dexamethasone solution (50 μM (w/v); Sigma Aldrich, United States) was added to the media on days 2 and 7. Shoot development was monitored and the number of shoots was recorded 40 days after transformation.

### RNA extraction, cDNA synthesis, and real time-PCR analysis

Tobacco leaf discs were excised and placed onto SIM and samples were harvested at 0, 3, 6, 9, 12, 15, and 18 days for RNA extraction. Total RNA was isolated from ∼100 mg leaf tissue samples using TRIZOL reagent (Sigma Life Science, USA). The quality and quantity of the RNA samples were analysed by agarose gel electrophoresis and Nanodrop spectrophotometer (Thermo Fisher Scientific, Waltham, Massachusetts, United States). High-quality total RNA samples with an A260/A280 ratio between 1.8 and 2, A260/A230 ratio between 2.0 and 2.4 were used for cDNA synthesis, using 1 μg of total RNA and the Verso cDNA synthesis kit (Thermo Fisher Scientific, United States). The synthesized cDNAs were then diluted 10 times (V/V, 1:10), and 2 μl of this diluted cDNA were used to amplify the target genes using the primers listed in the **Supplementary Table 2**. The *ACTIN* gene of tobacco (Genbank: GQ339768) was used as an internal control for normalization. For qRT-PCR analysis, 10 μl reaction mixture contained 2 μl of cDNA, 1 μl of each primer (from 5 μM stock), 5 μl SYBR green (Agilent Technologies, CA, United States), and 2 μl nuclease-free water. The reaction was carried out in Rotor-Gene 6000 real-time PCR machine (Corbett Life Science, Mortlake NSW 2137, Australia) using the following thermal cycling conditions: initial denaturation at 95 °C for 2 min, followed by 40 cycles at 95 °C for 15 s, 60 °C for 20 s, and 72 °C for 30 s. All the reactions were performed in biological triplicate. The fold-change in the expression of transcripts was calculated using the standard 2^−ΔΔCT^ method (Livak and Schmittgen, 2001). The expression patterns of the transcripts were plotted using Microsoft Excel 2007.

### Statistical Analyses

Confocal images presented throughout the study represent at least ten independent samples. All the experiments were conducted using three biological replicates. Significant differences among means were calculated at the 5% (*P* < 0.05) level, using Tukey’s HSD test through JMP software version 15 SAS Institute Inc., Cary, NC, 1989–2021.

## Acknowledgments

We are very thankful to Tzahi Arazi and Alexander Goldschmidt for their precious advice and critical reading of this manuscript. We are also grateful to Dr. Idan Efroni for providing the *TCSn/DR5* construct. M.K. would like to thank The Agricultural Research Organization (ARO), Volcani Center, Israel, for supporting this work with the ARO postdoctoral fellowships.

## Authors’ contributions

All the authors read and approved the final manuscript.

## Supplementary information

The online version contains supplementary material available at

## Data availability

The data underlying this article are available in the article and in its online supplementary material.

## Supplementary Tables

**Supplementary Table 1:** Media used to induce morphogenesis in tobacco explants.

**Supplementary Table 2:** Primers used in this study.

**Supplementary Table 3:** Details of reporter lines used in this study.

## Supplementary Figures

**Supplementary Fig. 1:** Developmental response of tobacco leaves placed on SIM or RIM. Bright light images of tobacco leaf explants exposed to SIM or RIM for 10, 14, 21 and 30 days.

**Supplementary Fig. 2:** Scanning electron microscope (SEM) analysis of tobacco leaf explants exposed to SIM and RIM, and relative transcript level of *NtCLV1* in tobacco explants exposed to SIM.. SEM images of tobacco leaf explants exposed to SIM (**a**) or RIM (**b**) for 7, 9, 11, 14, and 18 days. Scale bar: 1 mm.

**Supplementary Fig. 3:** Analysis of *proNtML1* activity in fully grown meristem and during *de novo* SAM formation in tobacco leaf explants.

**a.** Schematic representation of the T-DNAs built for this assay, consisting of the *mNG*, and *RFP* genes placed under the control of the *proNtML1* and *proAtUbiq*, respectively.
**b.** Confocal images of shoots excised from T0 transgenic tobacco plants obtained with the abovementioned construct. Scale bar: 50 μm.
**c.** Confocal images of transgenic T1 tobacco leaf discs exposed to SIM. Scale bar: 10 μm.

**Supplementary Fig. 4:** Relative transcript level of the *mSC* (**a**) and *mNG* (**b**) genes in transgenic tobacco leaf explants transformed with the construct presented in **Fig. 1d**, exposed to SIM for 0, 3, 6, 9, 12, 15, and 18 days. The data represent the mean ± SD from three biological replicates, n=3.

**Supplementary Fig. 5:** Analysis of *proNtWUS* and *proNtML1* activity during *de novo* SAM formation in tobacco plants.

**a.** Schematic representation of the T-DNAs built for this assay, consisting of the *mSC*, and *mNG* genes placed under the control of the *proNtWUS,* and pro*NtML1* respectively.
**b.** Confocal images of transgenic tobacco leaf discs exposed to SIM for 7, 12, 14, and 21 days. Scale bar: 20 μm.

**Supplementary Fig. 6:** Analysis of *NtML1* expression patterns during *de novo* SAM formation in tobacco leaf explants.

**a.** Schematic representation of the T-DNAs built for this assay, consisting of the *mNG*, and *RFP* genes placed under the control of the pro*NtML1* and *proAtUbiq*, respectively.
**b.** Confocal imaging of transgenic tobacco leaf discs exposed to SIM for 21 days. Scale bar: 20 μm

**Supplementary Fig. 7:** Alterations of L1 identity blocks SAM formation.

**a.** Schematic representation of the T-DNAs built for this assay, consisting of *proML1*-directed RNAi cassettes designed to target *NtML1* in L1, and the *mNG*, and *RFP* genes placed under the control of the pro*NtML1* and *proAtUbiq*, respectively.
**b.** Bright light and fluorescence images of tobacco leaf discs transformed with the abovementioned construct and exposed to SIM for 35 days. Scale bar: 500 μm
**c.** SEM images of tobacco explants transformed with the abovementioned constructs and exposed to SIM for 21 days. Scale bar: 1 mm
**d.** Histological images of tobacco explants transformed with the abovementioned constructs and exposed to SIM for 21 days. Scale bars: 50 and 100 μm.
**e.** Confocal imaging of tobacco explants transformed with the abovementioned constructs and exposed to SIM for 21 days. Scale bar: 20 μm.

**Supplementary Fig. 8:** *NtML1* overexpression blocks SAM formation.

**a.** Schematic representation of the T-DNAs built for this assay, consisting of *proSlUbiq*-directed *NtML1*overexpression cassette, and the *mNG* and *RFP* genes placed under the control of the pro*NtML1* and *proAtUbiq*, respectively.
**b.** Bright light images of tobacco explants transformed with the abovementioned constructs and exposed to SIM for 35 days. Scale bar: 500 μm
**c.** SEM images of tobacco explants transformed with the abovementioned constructs T-DNAsand exposed to SIM for 21 days. Scale bar: 1 mm
**d.** Confocal imaging of tobacco explants transformed with the abovementioned constructs and exposed to SIM for 21 days. Scale bar: 20 μm

**Supplementary Fig. 9:** DTA-mediated L1 layer ablation blocks early events of SAM formation.

**a.** Schematic representation of the T-DNAs built for this assay, consisting of the *mNG*, and *RFP* genes placed under the control of the pro*NtML1* and *proAtUbiq*, respectively, and of *DTA* directed by the *proSlUbiq* or *proNtML1*.
**b.** Bright light images of tobacco explants transformed with the abovementioned constructs and exposed to SIM for 35 days.
**c.** SEM images of tobacco explants transformed with the abovementioned constructs T-DNAs and exposed to SIM for 21 days. Scale bar: 500 μm.
**d.** Bright light and fluorescence images of tobacco explants transformed with the abovementioned constructs and exposed to SIM for 35 days.
**e.** Confocal imaging of tobacco explants transformed with the abovementioned constructs and exposed to SIM for 21 days. Scale bar: 100 μm.

**Supplementary Fig. 10:** Silencing *NtLOG4* blocks L1 identity and SAM formation.

**a.** Schematic representation of the T-DNAs built for this assay, consisting of the *proML1*-directed RNAi cassette designed to target *NtLOG4*, and the *mNG*, and *RFP* genes placed under the control of the *proNtML1* and *proAtUbiq*, respectively.
**b.** Bright light and fluorescence images of tobacco explants transformed with the abovementioned constructs and exposed to SIM for 35 days. Scale bar: 500 μm
**c.** SEM images of tobacco explants transformed with the abovementioned constructs T-DNAs and exposed to SIM for 21 days. Scale bar: 1mm.
**d.** Confocal imaging of tobacco explants transformed with the abovementioned constructs and exposed to SIM for 21 days. Scale bar: 20 μm.

**Supplementary Fig. 11**: Hormone signaling is active in L1 during tobacco regeneration.

**a.** Schematic representation of the T-DNAs built for this assay consisting of the mNG and mSC markers placed under the control of the *TCSv2* and *DR5* promoters.
**b.** Z-stack confocal imaging of tobacco explants transformed with the abovementioned constructs and exposed to SIM for 5, 7, 14 and 21 days. Scale bar: 20 μm.

**Supplementary Fig. 12:** Alteration of the process that directs anticlinal divisions blocks L1 and SAM formation.

**a.** Schematic representation of the T-DNAs built for this assay, consisting of *proML1*- and *proSlUBIQ*-directed RNAi cassettes designed to target *NtTON1*, and the *mNG*, and *RFP* genes placed under the control of the pro*NtML1* and *proAtUbiq*, respectively.
**b.** Bright light images of tobacco explants transformed with the abovementioned construct and exposed to SIM for 21 days. Scale bars: 500 μm (upper panel) and 1 cm (below panel).

**Supplementary Fig. 13:** Alterations of meristem functions block early events of SAM formation.

**a.** Schematic representation of the T-DNAs built for this assay, consisting of *proSlUBIQ*-directed RNAi cassettes designed to target *NtCLV1, NtPIN1*, and *NtHAM1*, and of the *mNG* and *RFP* genes placed under the control of the pro*NtML1* and *proAtUbiq*, respectively.
**b.** Bright light and fluorescence images of tobacco explants transformed with the abovementioned constructs and exposed to SIM for 35 days. Scale bar: 500 μm.
**c.** Bar graph depicting the number of developing shoots recorded 50 DAT with the abovementioned T-DNAs. The data represent the mean ± SD from three biological replicates, n=3; Student’s *t* test: \**P* < 0.05; \*\**P* < 0.01.
**d.** Confocal imaging of tobacco explants transformed with the abovementioned constructs and exposed to SIM for 21 days. Scale bar: 20 μm.

